# High quality, high throughput, and low-cost simultaneous video recording of 60 animals in operant chambers using PiRATeMC

**DOI:** 10.1101/2023.11.13.566747

**Authors:** Jarryd Ramborger, Sumay Kalra, Alexander C.W. Smith, Olivier George

## Abstract

**Background:** The development of Raspberry Pi-based recording devices for video analyses of drug self-administration studies has shown to be promising in terms of affordability, customizability, and capacity to extract in-depth behavioral patterns. Yet, most video recording systems are limited to a few cameras making them incompatible with large-scale studies.

**New Method:** We expanded the PiRATeMC (Pi-based Remote Acquisition Technology for Motion Capture) recording system by increasing its scale, modifying its code, and adding equipment to accommodate large-scale video acquisition, accompanied by data on the throughput capabilities, video fidelity, synchronicity of devices, and comparisons between the Raspberry Pi 3B+ and 4B models.

**Results:** Using PiRATeMC default recording parameters resulted in minimal storage (∼350MB/h), high throughput (< ∼120 seconds/Pi), high video fidelity, and synchronicity within ∼0.02 seconds, affording the ability to simultaneously record 60 animals in individual self-administration chambers at a fraction of current commercial costs. No consequential differences were found between Raspberry Pi 3B+ and 4B models.

**Comparison with Existing Method(s):** This system allows greater acquisition of video data simultaneously than other video recording systems by an order of magnitude with less storage needs and lower costs. Additionally, we report in-depth quantitative assessments of throughput, fidelity, and synchronicity, displaying real-time system capabilities.

**Conclusions:** The system presented is able to be fully installed in a month’s time by a single technician and provides a scalable, low cost, and quality-assured procedure with a high-degree of customization and synchronicity between recording devices, capable of recording a large number of subjects with high turnover in a variety of species and settings.

## 1. Introduction

The use of animal models in operant self-administration paradigms is critical to the advancements in the characterization of addiction behaviors, the mapping of neurobiological mechanisms, and their translation into human treatments and interventions (Spanagel, 2017). Animal models of drug self-administration typically use variables such as the number of infusions, active lever press, resistance to punishment, or breaking point to measure psychological construct such as “craving”, “motivation”, “escalation”, and “vulnerability” (Belin et al. 2009; Guglielmo et al. 2023, preprint, doi:10.1101/2021.07.22.453410; Kallupi et al. 2022, preprint, doi:10.1101/2022.07.26.501618). However, these simple behavioral measures are limited in their capability to provide in-depth information about the organization and the complexity of behavior during an operant session. The advancements in machine-learning-based video analyses provides an opportunity to identify behavioral motifs, complex behavioral changes, and predict behaviors of interest (Mathis et al., 2018; Nath et al., 2019; Luxem et al., 2022; Pereira et al., 2022). Current video recording systems are either expensive proprietary technologies (MedAssociates, The Imaging Source, GoPro, Point Grey, Basler, etc.) or inexpensive open-source systems (usually based on Arduino and Raspberry Pi) that currently lack demonstrated scalability, with most systems validating no more than 1-16 cameras recorded simultaneously in a variety of species and systems (Saxena et al., 2018; Singh et al., 2019; Weber and Fisher, 2019, preprint, doi:10.1101/596106; Hou and Glover, 2022; Marcus et al., 2022; Centanni and Smith, 2023).

To address this gap in the literature, we tested the capability, throughput, fidelity, and synchronicity of videos of the PiRATeMC (Pi-based Remote Acquisition Technology for Motion Capture) protocol by Centanni and Smith (2023). This study also provides modifications of the code to simultaneously record 60 animals in independent operant chambers, along with functional comparisons between models of Raspberry Pis, and system comparisons with other Pi-based and commercially-available alternatives. A link to a GitHub providing a tutorial in greater detail for transparency and replication of system design, as well as the code block for FFmpeg tests and analyses, is also provided.

## 2. Methods and Materials

Throughout the present paper, any file referenced in the PiRATeMC system will be in *italics*, while any text that is a code typed into a terminal of a computer or the Raspberry Pis (RPi) will utilize the same convention as Centanni and Smith (2023) by ***bolding and italicizing*** as such: ***terminal_code*.**

### 2.1. Assembly and Installation

#### 2.1.1. PiRATeMC System

The PiRATeMC (Pi-based Remote Acquisition Technology for Motion Capture) system as outlined in Centanni and Smith (2023) for a cluster design was implemented. This includes the basic components, such as the RPis and their accessories, the network switch, and remote control, as well as the codes necessary to allow these three devices within the system to interface. An accessible instructional manual may be found on their GitHub (https://github.com/alexcwsmith/PiRATeMC/blob/master/README.md). In short, this requires assembly of the RPi parts, flashing a micro-SD card with an operating system image, installation of the latest Ubuntu operating system (OS) to the remote controller, lines of code for packages to be installed and files to be copied within the remote controller to interface with the network switch, connecting to the RPi and updating the recording file with your specifications, updating variables in the *.bashrc* of the RPi with remote controller information, updating the *.bashrc* file of the remote controller with the IP addresses to all RPis on its network, and running ***cssh*** with all RPi IP addresses in a terminal to interface with each connected RPi.

#### 2.1.2. Coding Alterations and Equipment Additions

In terms of coding, the call variables ***$REMOTEPASS*** and ***$REMOTEVIDPATH*** present in the *.bashrc* and *recordVideo.sh* files were not utilized due to the reoccurring errors ***permission denied***, referring to ***$REMOTEPASS*** calling the remote’s password, and ***no such directory*** referring to ***$REMOTEVIDPATH*** calling the path to the video storage location on the remote. To aid technicians in troubleshooting, ***echo*** components were added after each code step in *recordVideo.sh* to better locate coding errors. Lastly, to accommodate different experimental designs occurring simultaneously the *recordVideo.sh* file was saved as two separate codes with their respective video paths. A tutorial on how to install and configure the updated PiRATeMC system can be found on GitHub (https://github.com/George-LabX/raspicluster/blob/main/README.md). In terms of equipment, additions were the Arducam fisheye lens, Raspberry Pi PoE HAT+, Seagate 5TB external hard drives, and a locally manufactured acrylic device for secure camera installation to a modular chamber.

#### 2.1.3. Operant Chambers

The RPis were housed in an ENV-007CT Med Associates modular operant chambers (53.34cm x 34.93cm x 27.31cm) designed for rodents equipped with an ENV-005 grid floor (28.6cm x 24.1cm). A comprehensive inventory of the individual products utilized in this design are provided in Table 1. below including those in the PiRATeMC paradigm.

**Table 1.**
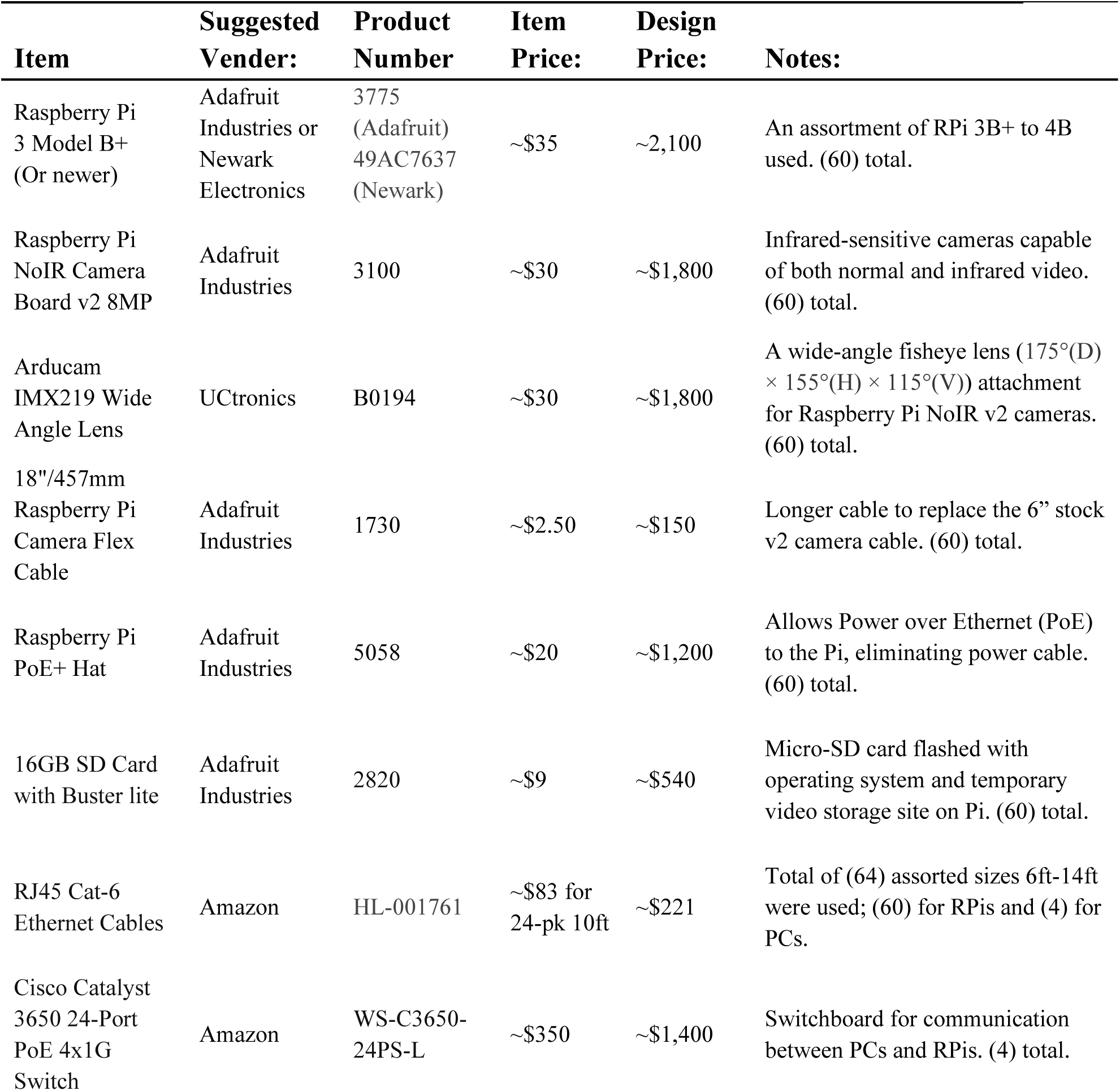
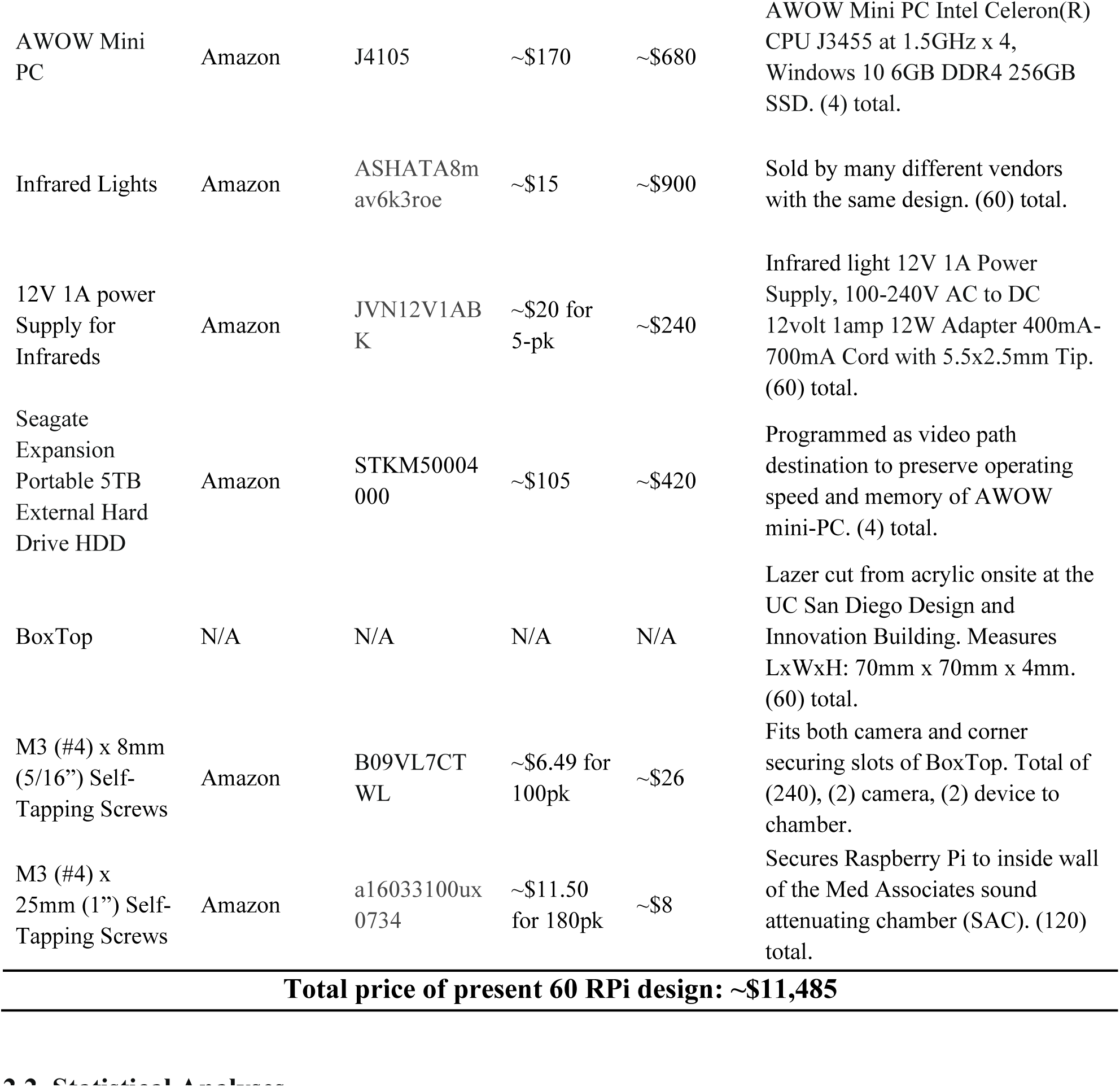
All products utilized including vendor, individual price, price within a 60 RPi design, and usage or variation notes, and a total cost.

### 2.2. Statistical Analyses

Data on video size was gathered by averaging 60 videos from a 1-hour mock session at the default parameters of 30 frames per second (FPS), 10M bitrate, and 800×600 resolution. To acquire data on throughput, fidelity, synchronicity, and functionality between RPi models, we ran four separate one and two hour sessions in mock conditions with 15 boxes for 2 hours (2h-15 Boxes), ten boxes for 2 hours (2h-10 Boxes), ten boxes for 1 hour (1h-10 Boxes), and five boxes for 1 hour (1h-5 Boxes) for scalable comparisons (Fig. 1.). Results of tests were analyzed using R Studio (R version 4.3.1 (2023-06-16 ucrt)). Preliminary analyses of QQ Plots, Shapiro-Wilk tests, and Levene’s tests were conducted to ensure the proper statistical comparisons were made.

**Fig. 1.**
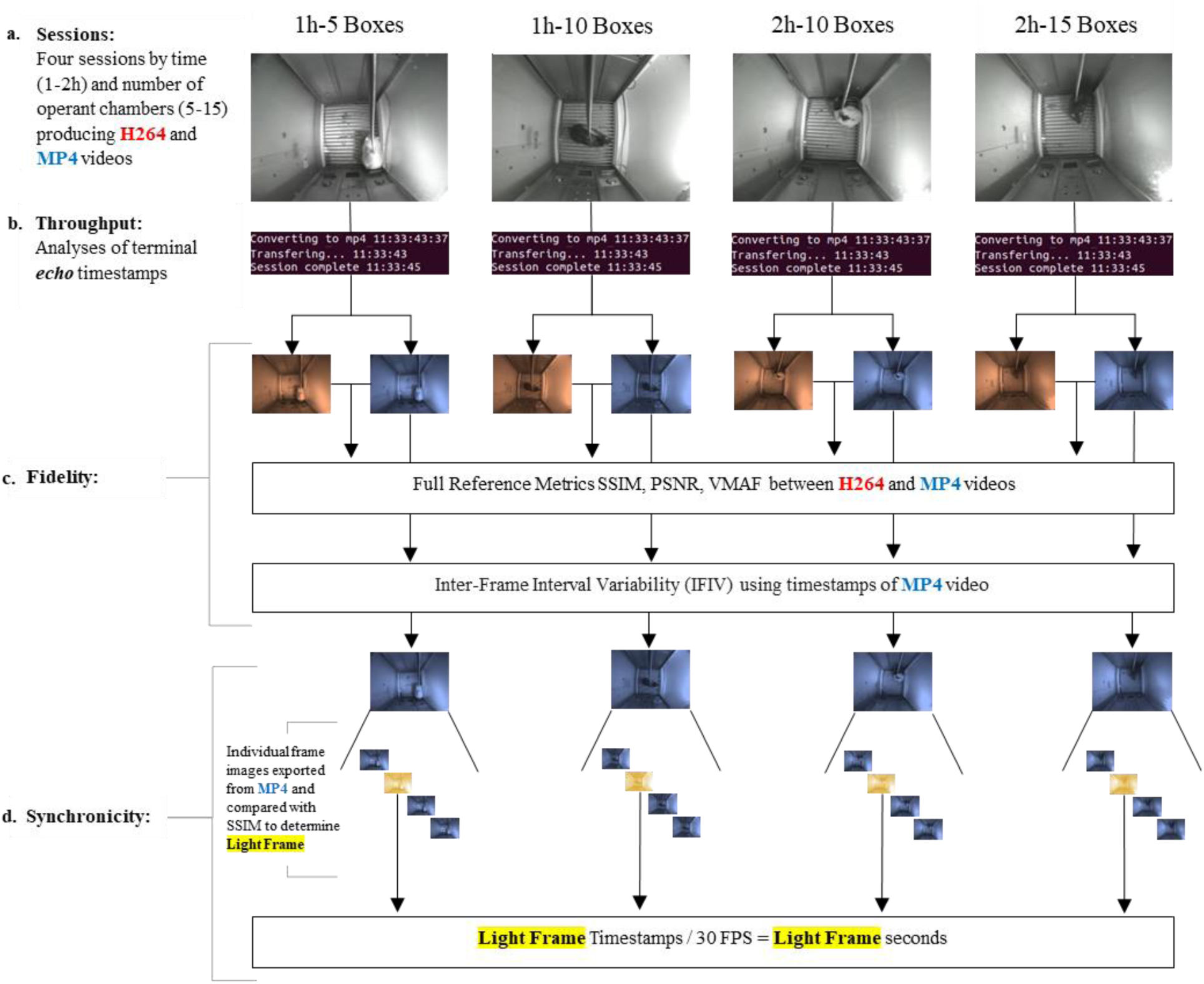
Workflow. Visualization of data acquisition and statistical analyses.

#### 2.2.1. Throughput

Throughput was analyzed by documenting the ***echo*** notations when the conversion to MP4 began (“Converting to mp4”), when the transfer to the remote began (“Transferring…”), and when the transfer was complete (“mp4 transfer complete”), modified in the PiRATeMC *recordVideo.sh* recording file for the four-session experiment. Conversion time (convert_secs) was defined as the time between the “Converting…” timestamp and the “Transferring…” timestamp, and transfer time (trans_secs) defined as the time between the “Transferring…” timestamp and the “…complete” timestamp. Medians, and standard error of each were subsequently computed as well as the full processing time (convert_secs + trans_secs = process_secs) from each RPi to remote PC (Fig. 1.b.).

#### 2.2.2. Fidelity

To investigate fidelity (Fig 1.c.), we modified *recordVideo.sh* not to delete the original H.264 file and to transfer it with the MP4 file to the remote PC. The H.264 (reference) video file and the distorted (product) MP4 video were analyzed with FFmpeg software in conjunction with Windows PowerShell using three full reference metrics (FR): Structural Similarity Index Measure (SSIM), Peak Signal-to-Noise Ratio (PSNR), and the Video Multimethod Assessment Fusion (VMAF). Separately, we computed the Inter-Frame Interval Variability (IFIV) for dropped frames.

SSIM is a measure of elements luminance, contrast, and structure that are relatively independent of one another and combined into a single score between -1 and 1 (Wang et al., 2004; Venkataramanan et al., 2021). PSNR is expressed as a decibel (dB) logarithm of the mean squared error (MSE) (Deshpande et al., 2018; Setiadi, 2021) of luma (Y) and chromatic (U, V) elements. The VMAF recently developed by *Netflix* is a combination of modified versions of the quality metrics Visual Information Fidelity (VIF) and Detail Loss Metric (DLM) in combination with a metric of motion computed from the luminance element (Li et al. 2016) and is found to be highly correlated with the human visual system (HSV).

SSIM, PSNR, and VMAF metrics of the system recordings were computed by exporting the results of each H.264 file and its corresponding MP4 file before computing the median and standard error of all videos. The IFIV was computed by exporting frame timestamps from each MP4 file, calculating the time between each timestamp, and the means, medians, and standard deviations were computed for each video and between all videos.

#### 2.2.3. Synchronicity

Synchronicity (Fig 1.d.) was determined by beginning a recording with all operant chambers closed and running a Med Associates program to initiate both cue lights for two seconds and determining for each RPi the moment the cue lights turned on and whether this was timestamped differently across all Pis. This was conducted by extracting the first 10 seconds in each video as an individual image and utilizing the SSIM analysis in FFmpeg. score. Utilizing a dark reference image and comparing it with all subsequently exported images at the peak of the light event, enables the identification of an inverse relationship between luminance (Y) and the event, with the lowest score representing the light event. Each RPi frame representing the light event was subsequently divided by the FPS (30) to get a time in seconds the event occurred. The mean and standard error was taken of all light frames.

#### 2.2.4. Categories and Variables

Recordings were compared by Time (1h and 2h) and by RPi Model (3B+ and 4B) to determine potential differences in length of session time and the most recent available Raspberry Pi models. Table 2. below displays the variables compared across the three categories complete with name, definition, and operationalization.

**Table 2.**
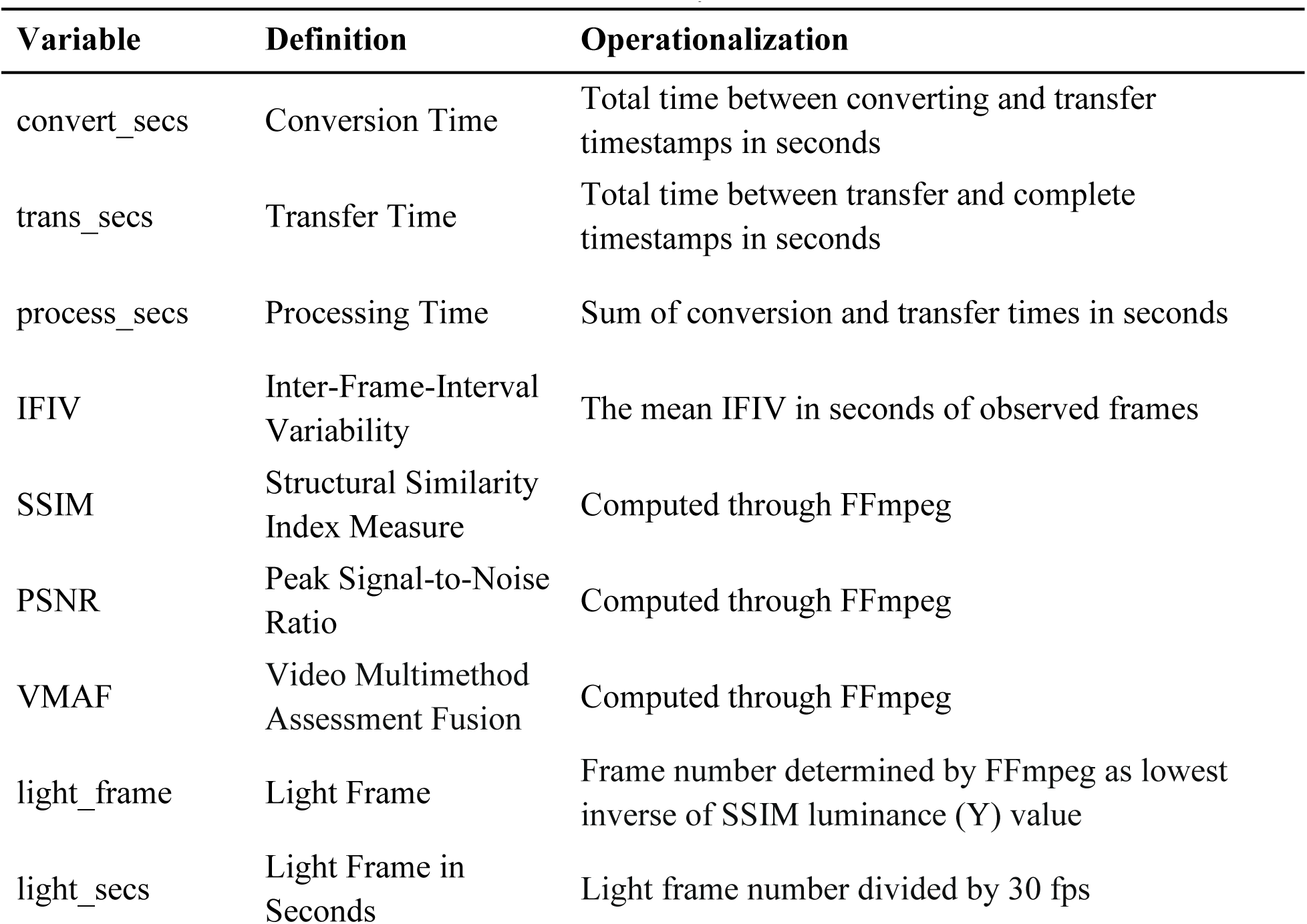
Table of variables used for statistical analyses

Normality and homogeneity of variances were tested using QQ Plots, Shapiro-Wilk tests, and Levene’s tests. The Time comparisons were conducted with Wilcoxon-Mann-Whitney tests due to non-normal distributions. All comparisons between RPi models for variables conversion time, processing time, IFIV, SSIM, PSNR, and VMAF were made with Wilcoxon-Mann-Whitney tests due to non-normal distributions, while transfer seconds was compared by Student’s t-test. The variables light frame and light seconds for all comparisons between groups were conducted by Levene’s test of variance given the recordings were conducted on separate computers and therefore tests of internal variability were preferred to determine synchronicity.

## 3. Results

### 3.1. Hardware Assembly

#### 3.1.1. Raspberry Pi and Camera Assembly and Installation

Figure 2. displays the parts necessary to assemble the system as instructed by Centanni and Smith (2023) with additions of the Arducam wide-angle fisheye lens ((175°(D)×155°(H)×115°(V)) (Fig. 2.c.) as the original NoIR v2 camera did not allow visibility of the chamber levers when secured to the top pane of glass (Fig. 3.). The RPis were then positioned on the inside wall of the outer sound attenuated box using two standard M3 x 1” self-tapping screws on the right wall at a distance corresponding to the width of the RPi circuit board. The RPis were secured in place with loose zip-ties to the screws by using the PoE+ HAT posts, allowing for easy removal or placement of the RPi and access to the micro-SD cards as needed (Fig. 7.a). This arrangement ensured that the RPis were in a location with sufficient airflow for cooling and allowed unobstructed access for the camera cable to reach its designated operating position, while also remaining close to the hole in the side of the box for the ethernet cable.

**Fig. 2.**
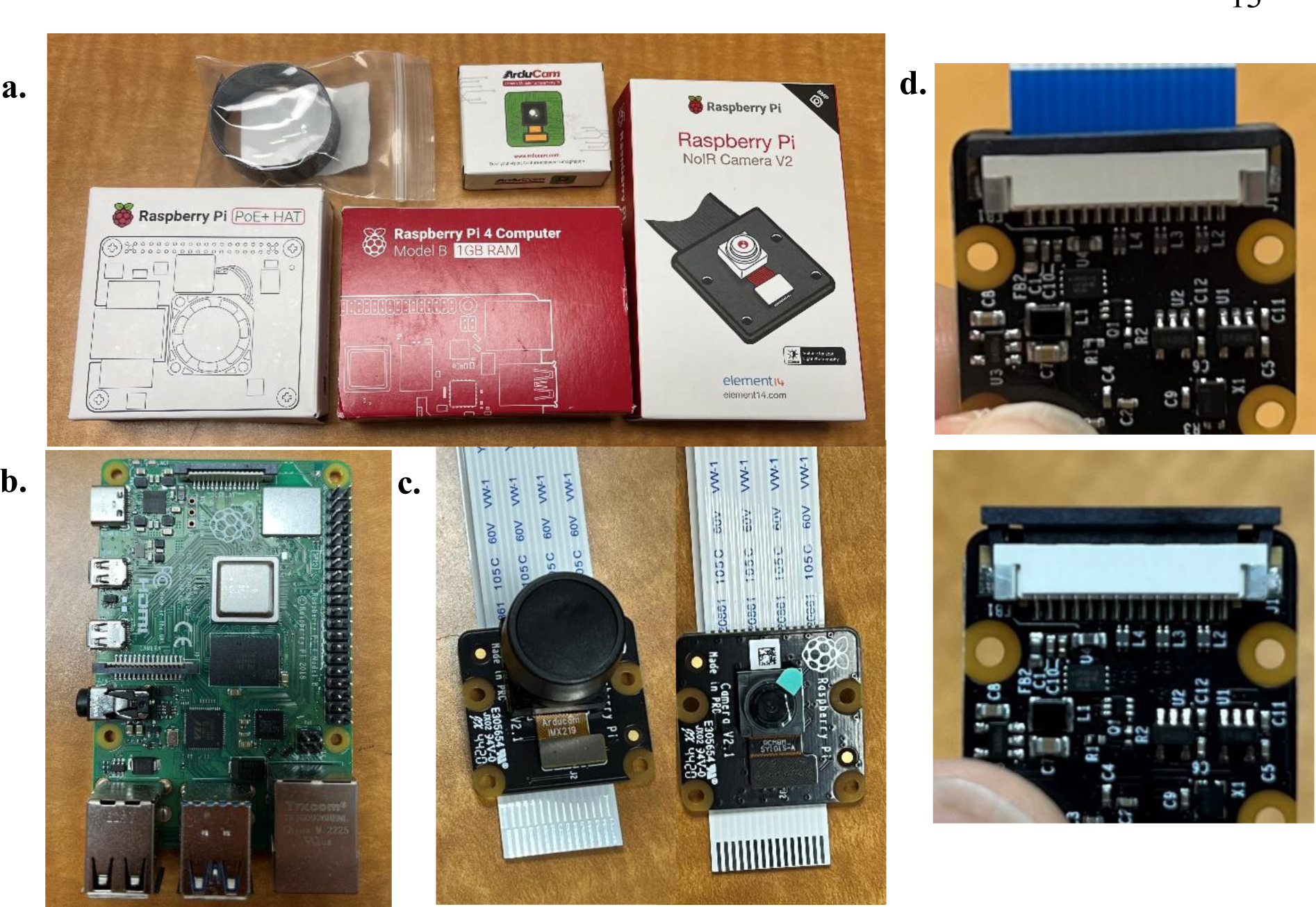
Raspberry Pi Assembly. **a.** Visualization of each individual Raspberry Pi product purchased, assembled, and utilized in this experiment. **b.** Unboxed Model 4B RPi with ports towards user. **c.** Fisheye lens (left) replacement for stock lens (right). **d.** Camera tabs to replace and install ribbon cable.

**Fig. 3.**
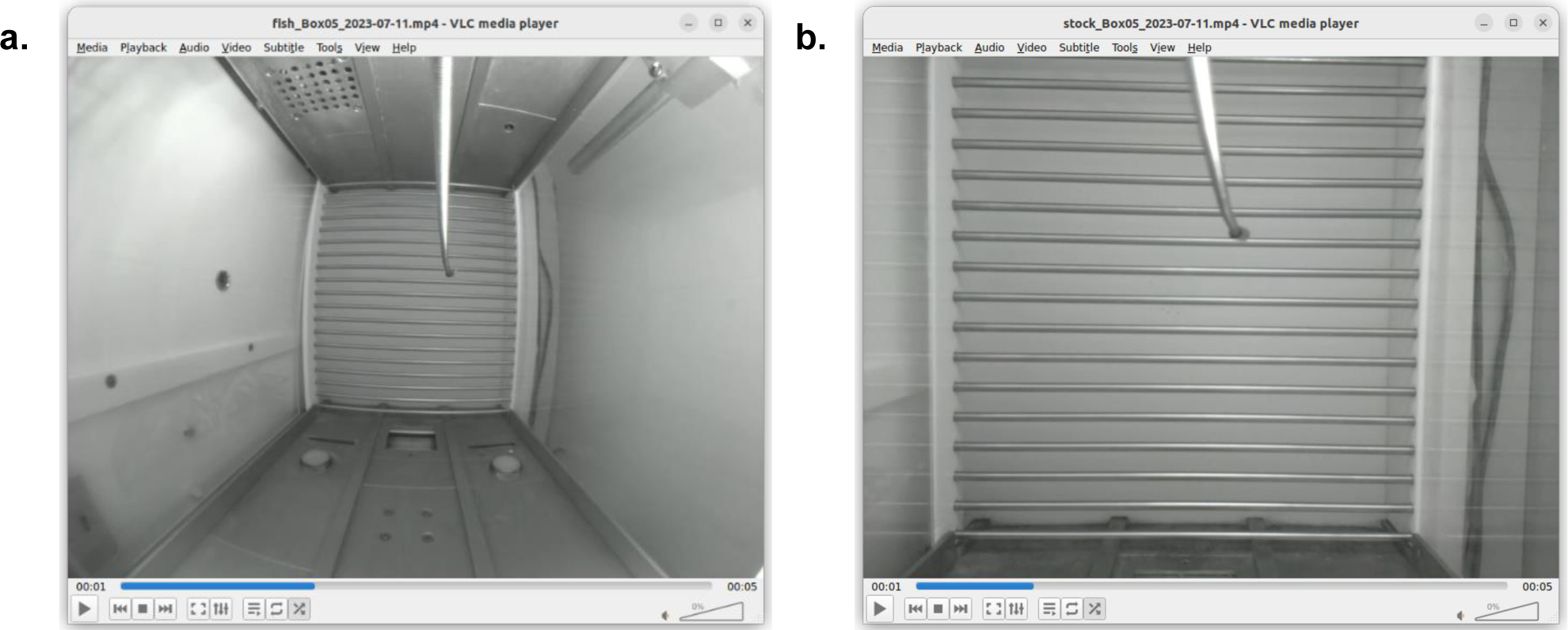
Visualization of camera comparisons. **a.** Arducam fisheye lens **b.** stock NoIR v2 camera. Both set at a resolution of 800×600.

To provide consistent recordings, the cameras need to be positioned in a location that has full visibility into the operant chamber without being obscured by the outer glass, not obstruct the catheter line into the chamber or the technician from installing the subject, nor be moved at any point during the experiment. To address these requirements, the cameras were secured to a device laser cut from acrylic designed specifically for this configuration called a BoxTop with sections removed for the catheter line and camera installation. This was achieved with two M3 x 8mm self-tapping screws through both the camera board and device. The top of the ENV-007CT modular chamber had a 5cm (50mm) diameter hole located at the center of the plexiglass or metal material, the BoxTop measures 70mm x 70mm x 4mm and was secured to the top of the operant chamber by two M3 x 8mm self-tapping screws at diametric corners (Fig. 4. and Fig. 5.). The lens measured approximately 28cm from the grid floor after installation.

**Fig. 4.**
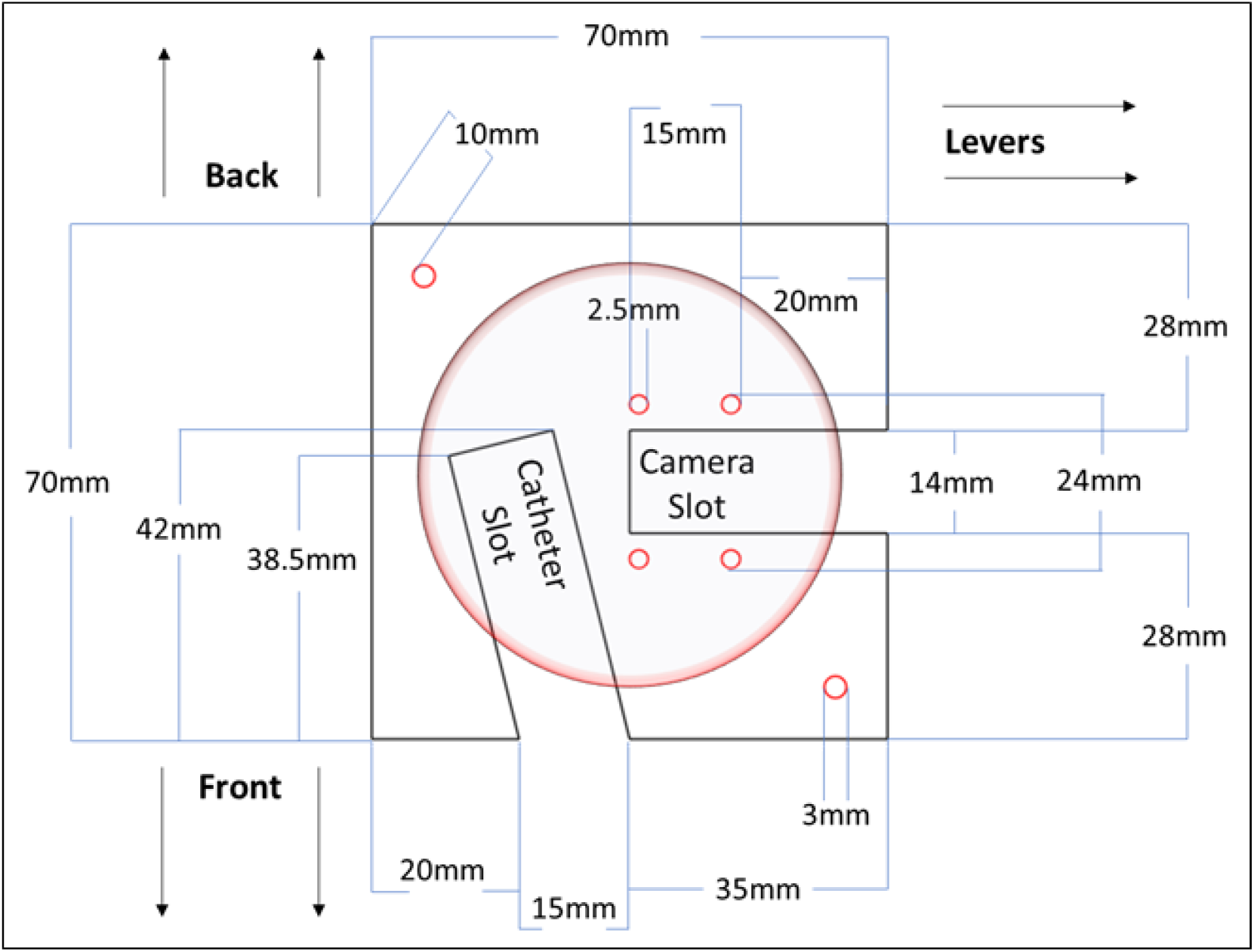
BoxTop. Visualization of the BoxTop schematic complete with measurements and display of placement over 50mm diameter operant chamber hole (large red center circle).

**Fig. 5.**
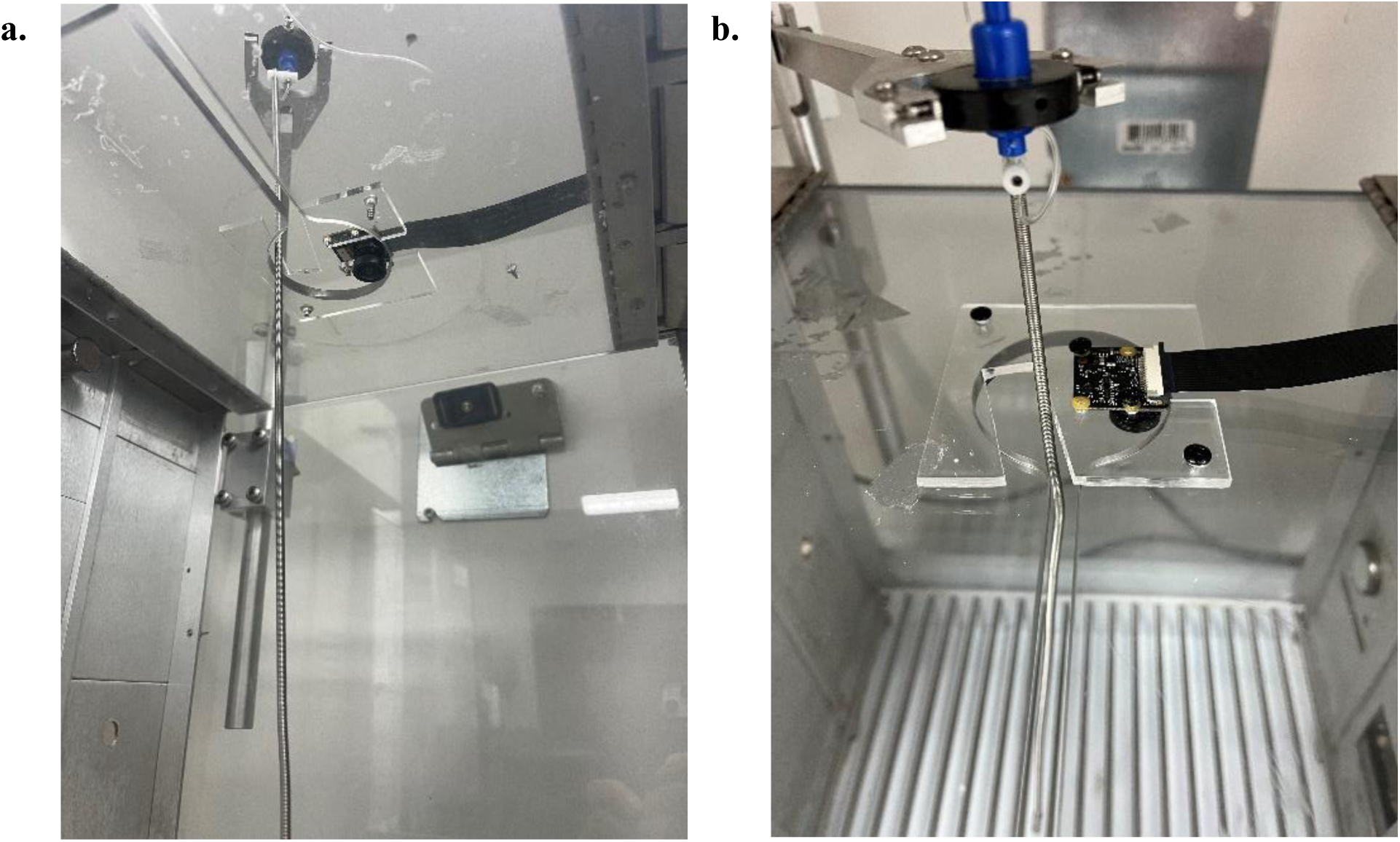
Visualization of the BoxTop installed inside an operant chamber. **a.** Subject’s eye-view from inside modular chamber. **b.** Technician view from top of chamber.

#### 3.1.2. Ethernet Switch

For multiple RPis to communicate over Ethernet, a centralized integration hub is needed. The Cisco 3650 Catalyst 24-port switch provides connection as well as power over Ethernet to the RPis thereby eliminating a power cable as well as simplicity in configuration with the remote controller. Approximately five minutes after being powered on the switch should be fully booted and ready to plug-and-play. This ethernet switch does not require internet connection and enables clustering and simultaneous communication with up to 23 RPis, while models are available with up to 48-ports, as well as methods to stack switches to accommodate ever increasing numbers of RPis on one network.

#### 3.1.3. Remote Controller

Now that a device for recording and a hub of integration is established, an instrument to dispatch commands in a synchronized fashion and receive the videos produced is necessary to function as a remote controller. The latest version of Ubuntu desktop is suggested to be installed to ensure seamless interaction with the Raspbian Linux operating system present on the RPis.

This is carried out with a USB drive containing at least 4GB of storage and creating a bootable Ubuntu image. A tutorial of the process can be found at https://ubuntu.com/tutorials/create-a-usb-stick-on-windows#1-overview. The AWOW mini-PC (and many other brands or PC options) can be found in multiple storage and RAM sizes to accommodate any individual preferences, while Fig. 6. shows the system information of those used in this design.

**Fig. 6.**
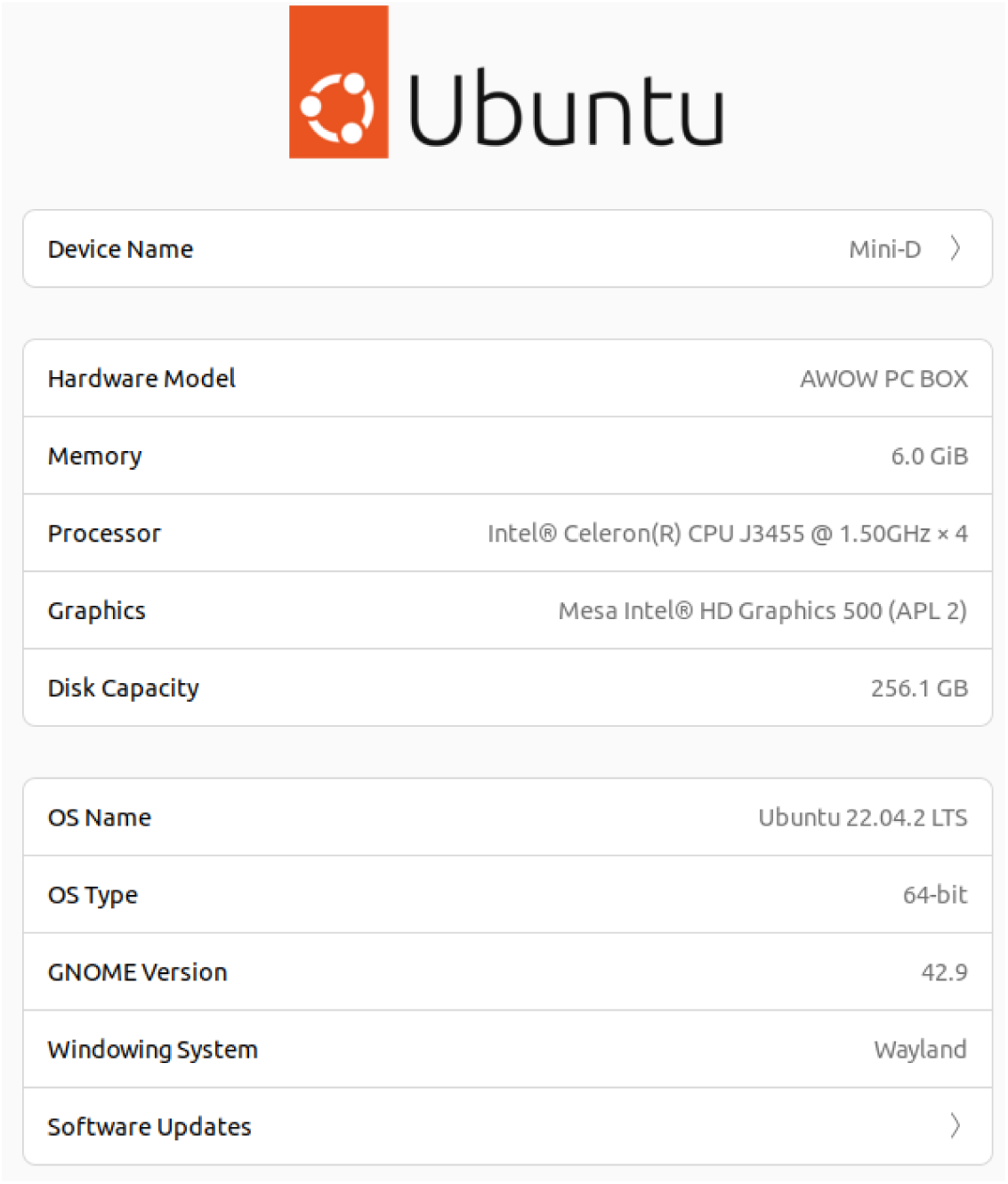
AWOW Mini-PC. Screen shot of AWOW Mini-PC system information.

#### 3.1.4. Video Path and Storage

Given the Linux operating system currently lacks the “online-only” feature of DropBox, each AWOW was fitted with a Seagate 5TB external hard drive in one of its two 3.0 USB ports for storage of recordings before being uploaded to DropBox via an internet browser.

#### 3.1.5. Infrared Installation

The infrared lights were installed approximately 1-2” from the ceiling on the back wall of the outer sound attenuating box (Fig. 7.b. and 7.c.), pointing directly into the chamber or downwards against the wall. The reflected light provides sufficient visibility for recordings or images. In the ENV-018V cubicles, which have a greater height clearance, lights were installed on the back right corner of the ENV-007CT modular operant chamber itself.

**Fig. 7.**
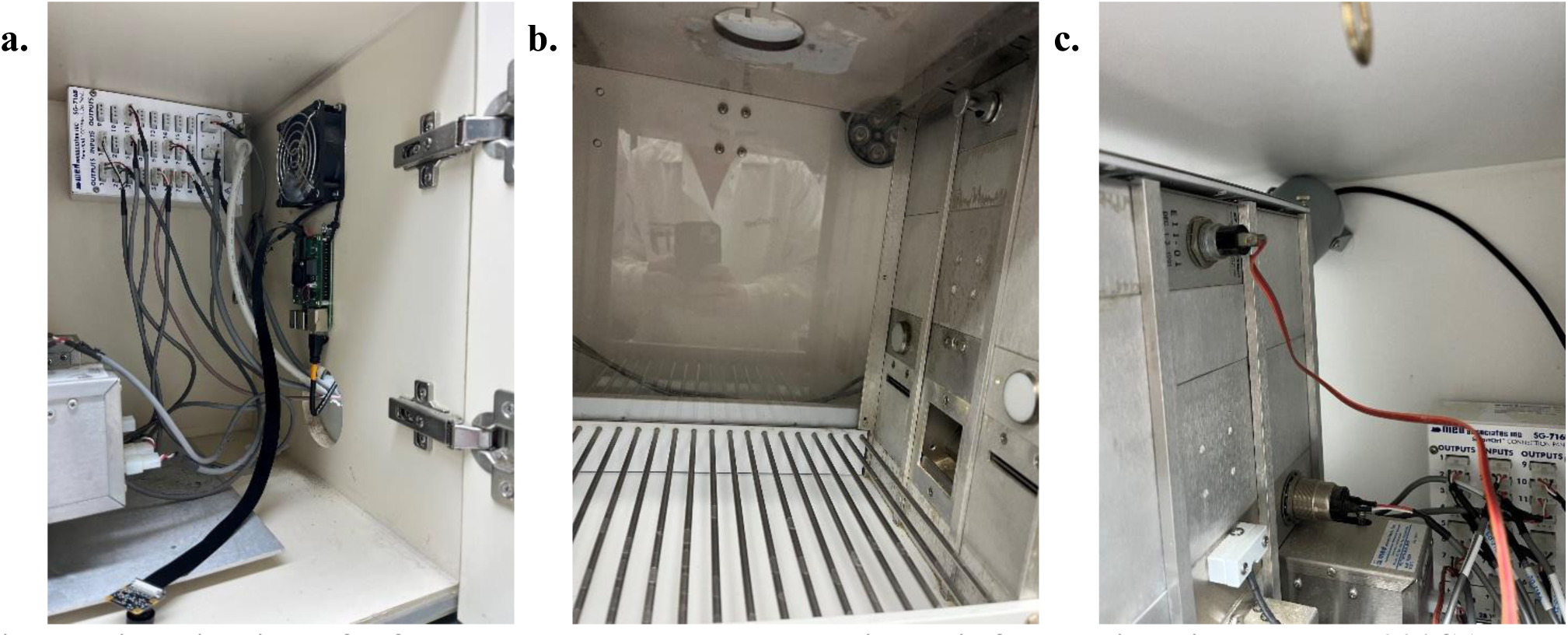
Visualization of a fully assembled Raspberry Pi and infrared light in an ENV-022SA Med Associate SAC. **a.** Raspberry Pi secured to inside wall by two M3 x 1” self-tapping screw and zip-ties. **b.** Inside view of infrared light secured to back wall of SAC by one M3 x 1” self-tapping screw pointing into chamber. **c.** Side view of infrared light secured to back wall of SAC.

### 3.2. Software Assembly and Functionality

#### 3.2.1. Additions and Remarks to *recordVideo.sh*

Recording multiple experiments containing different variables or identifiers can be challenging with a single recording file as it requires modifying the code on each Pi prior to the experiment. Furthermore, having a single location for all video files may become difficult to parse between the different experiments one may have running at any given time.

To accommodate this, separate video code files can be created for each variable or experiment identifier complete with their own respective video path to avoid the need to change the recording file for every different experiment that uses a different variable. This can be accomplished by editing the *recordVideo.sh* file with the desired path and parameters, then saving it with its respective name for identification and execution.

For example, *reccoc.sh* (“<u>rec</u>ord <u>coc</u>aine”, Fig.8.) is one created from the *recordVideo.sh* file complete with its respective path to its folder on the Seagate hard drive (Expansion). Once created, the file is then made executable with “***chmod +x***”, meaning, “change mode of file to allow execution”, in which case, ***chmod +x reccoc.sh***, in this example.

**Fig. 8.**
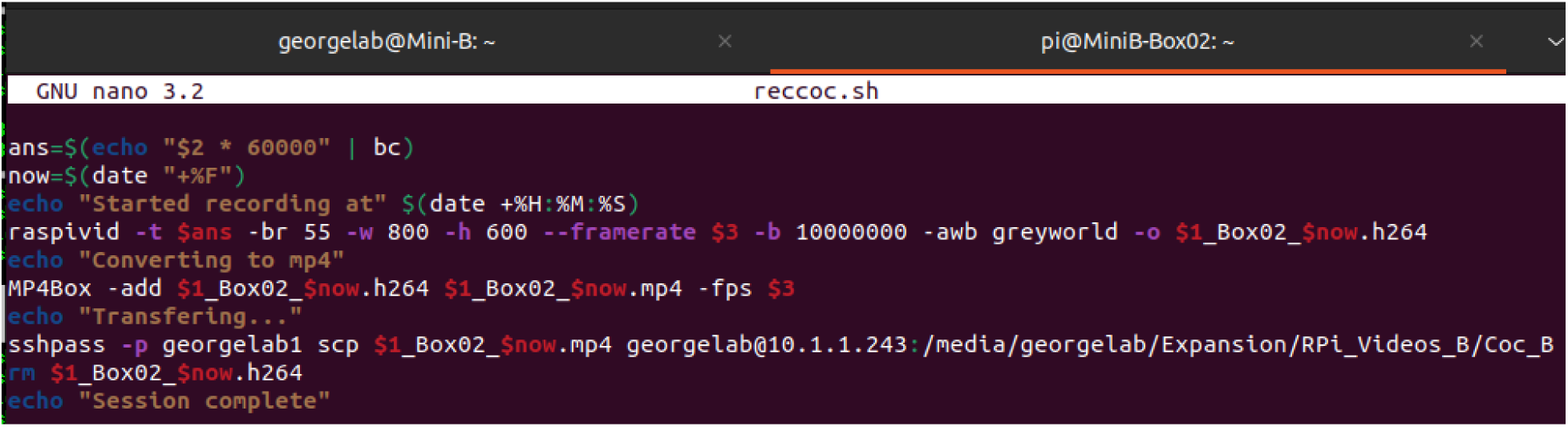
Example of the *reccoc.sh* executable file to record and transfer videos to the remote.

#### 3.2.2. Execution of *recordVideo.sh*

Once materials are installed, programming connections and interfaces effectively established, and recording parameters are set, execution of the *recordVideo.sh* file begins the recording function on all RPis connected to a switch simultaneously (Fig. 9.).

**Fig. 9.**
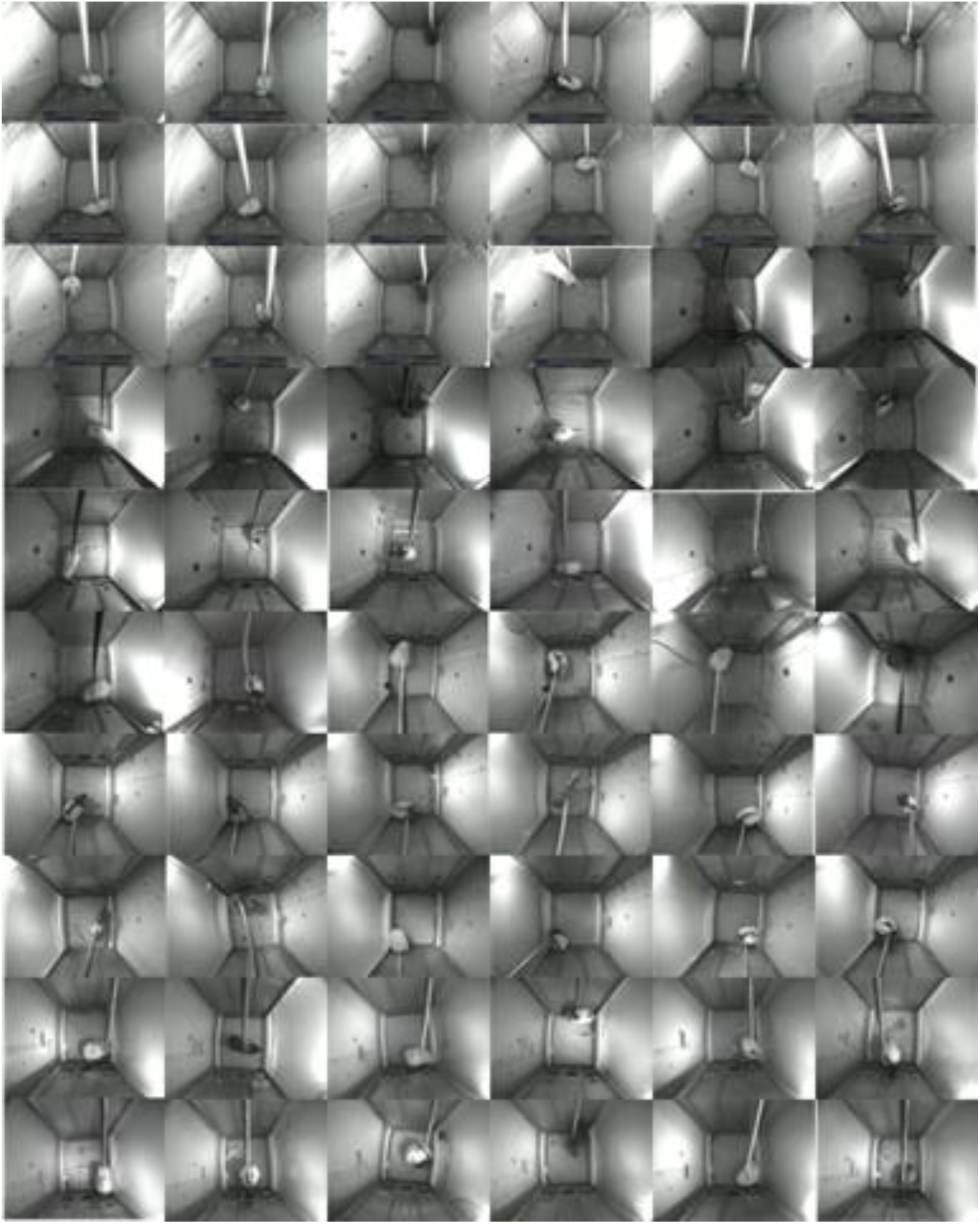
Recording of 60 Raspberry Pis simultaneously across four AWOW mini-PCs.

### 3.3. Raspberry Pi Quality Assessments

#### 3.3.1. Video Size

Commercial systems such as Med Associates do not allow customization of video parameters to suit individual needs in terms of field of view and storage capacities. Only two parameters in the *recordVideo.sh* file of the PiRATeMC paradigm affect the resulting video size: bitrate (set at 10M, raspivid defaults at 17M) and resolution (set at 800×600). When executing the *recordVideo.sh* file set at 30 frames per second, a bit rate of 10M, and 800×600 resolution, the average one-hour video is approximately ∼350MB per RPi. For the current system with 60 RPis that results in about ∼21GB per hour, ∼127GB for a 6-hour self-administration session, and ∼254GB for a 12-hour session.

If one were to establish a single AWOW and 24-port Catalyst design with the maximum 23 RPis, that equates to ∼8.13GB total of video every hour. Considering the micro-SD cards on the RPis are only 16GB (larger ones are available) and the average hour of video is ∼350MB the micro-SD cards will need to be cleared at most every ∼45 hours of video, while a 256GB model AWOW connected to 23 RPis will need to every ∼31 hours (not accounting for storage allocated to OS systems on each device).

#### 3.3.2. System Fidelity

Analyses across the system (N=40) for fidelity measures SSIM (0.99 ± 3e-4), VMAF (94 ± 0.2), and PSNR (40 ± 0.4), as well as IFIV (0.03 ± 3e-4) revealed high levels of quality between recordings, with coefficient of variances of 0.19% (SSIM), 1.11% (VMAF), and 5.67% (PSNR), while only one RPi had a single dropped frame.

#### 3.3.3. Throughput Comparisons By Time

Descriptive analyses of the four separate sessions showed that any given RPi (N=40), irrespective of number of RPis or length of session, had a median conversion time of 55 ± 12 seconds and a median transfer time of 48 ± 11 seconds, for a total median processing time of 116 ± 22 seconds.

Comparison analyses of throughput by Time were conducted with Wilcoxon-Mann-Whitney tests. Medians for throughput variables were 27 ± 10 seconds (1h) and 70 ± 16 seconds (2h) for conversion time, 32 ± 5 seconds (1h) and 50 ± 16 seconds (2h) for transfer time, and 48 ± 15 seconds (1h) and 129 ± 30 seconds (2h) for processing time, revealing an effect of scale as a 2h session required 2.7 times more processing time than a 1h session. A significant difference was found between the session lengths in the conversion time variable (*U*(15, 25) = 36.5, *p* = < .001) with a large effect size (*r* = 0.667) (Fig. 10.a.). Similar significance values and large effects were found in transfer time (*U*(15, 25) = 54.5, *p* = < .001, *r* = 0.588) and in processing time (*U*(15, 25) = 38.5, *p* = < .001, *r* = 0.659) between the 1h and 2h session lengths (Fig. 10.b. & 10.c.).

**Fig. 10.**
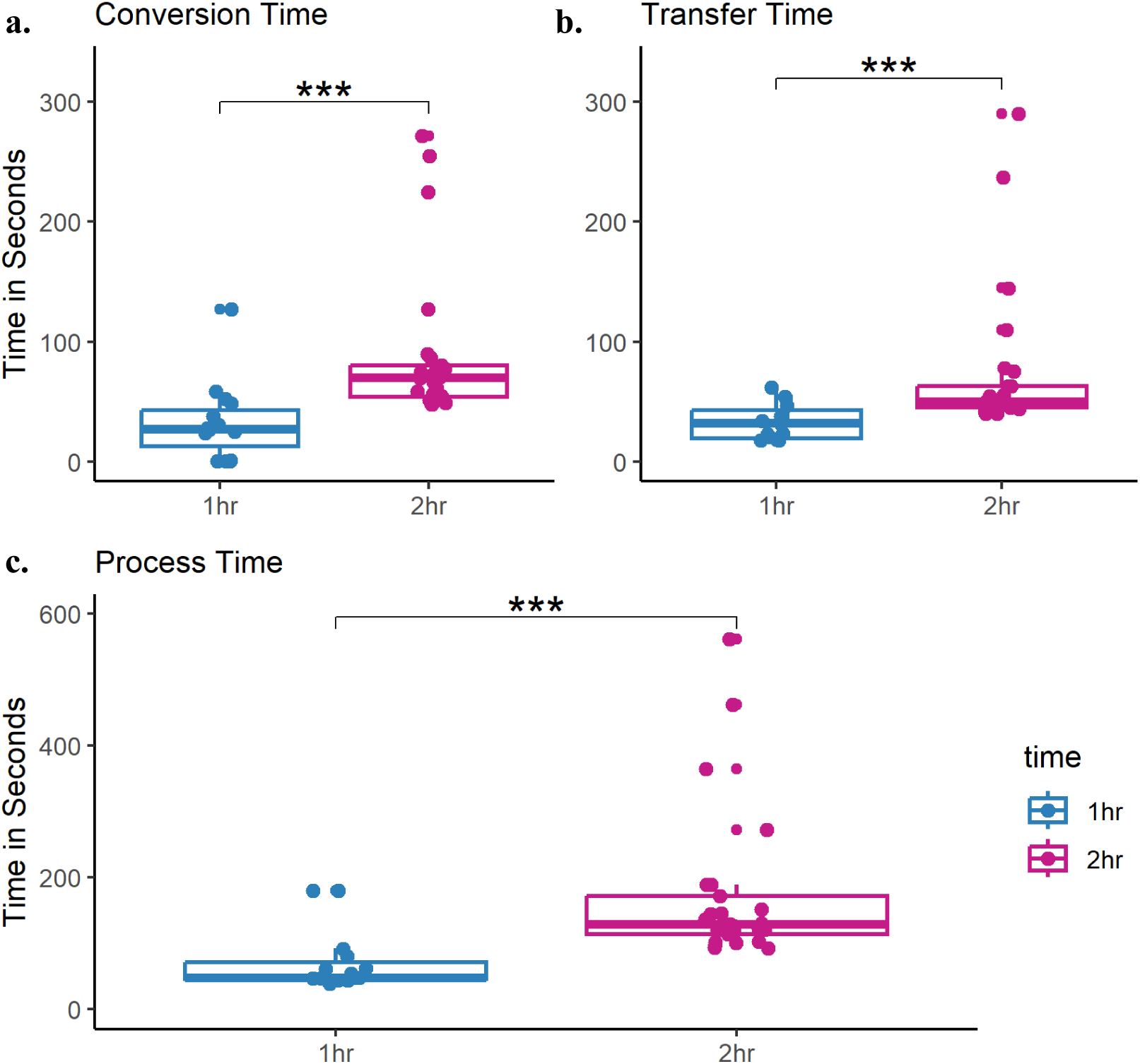
Boxplots of throughput comparisons between length of Session (Time). Conversion Time (a.), Transfer Time (b.), and Process Time (c.).

#### 3.3.4. Raspberry Pi Model 3B+ and 4B Comparisons

##### 3.3.4.1. Throughput

Throughput analyses grouped by Pi Model revealed median conversion times of 0 ± 0.3 seconds (3B+) and 30 ± 12 seconds (4B), mean transfer times of 50 ± 5 seconds (3B+) and 27 ± 9 seconds (4B), and median processing times of 51 ± 6 seconds (3B+) and 48 ± 23 seconds (4B). Wilcoxon-Mann-Whitney tests determined a significant difference in conversion time between the models (*U*(4, 11) = 0, *p* = .005) with a large effect size (*r* = 0.744) (Fig. 11.a.). Student’s t-test revealed a significant difference between model 3B+ (*M* = 50.25, *SD* = 10.21) and model 4B (*M* = 26.82, *SD* = 11.3) in transfer time (*t*(5.92) = 3.82, *p* = .009) (Fig. 11.b.), while no significant differences were found between models in processing time (*U*(4, 11) = 16.5, *p* = .511).

**Fig. 11.**
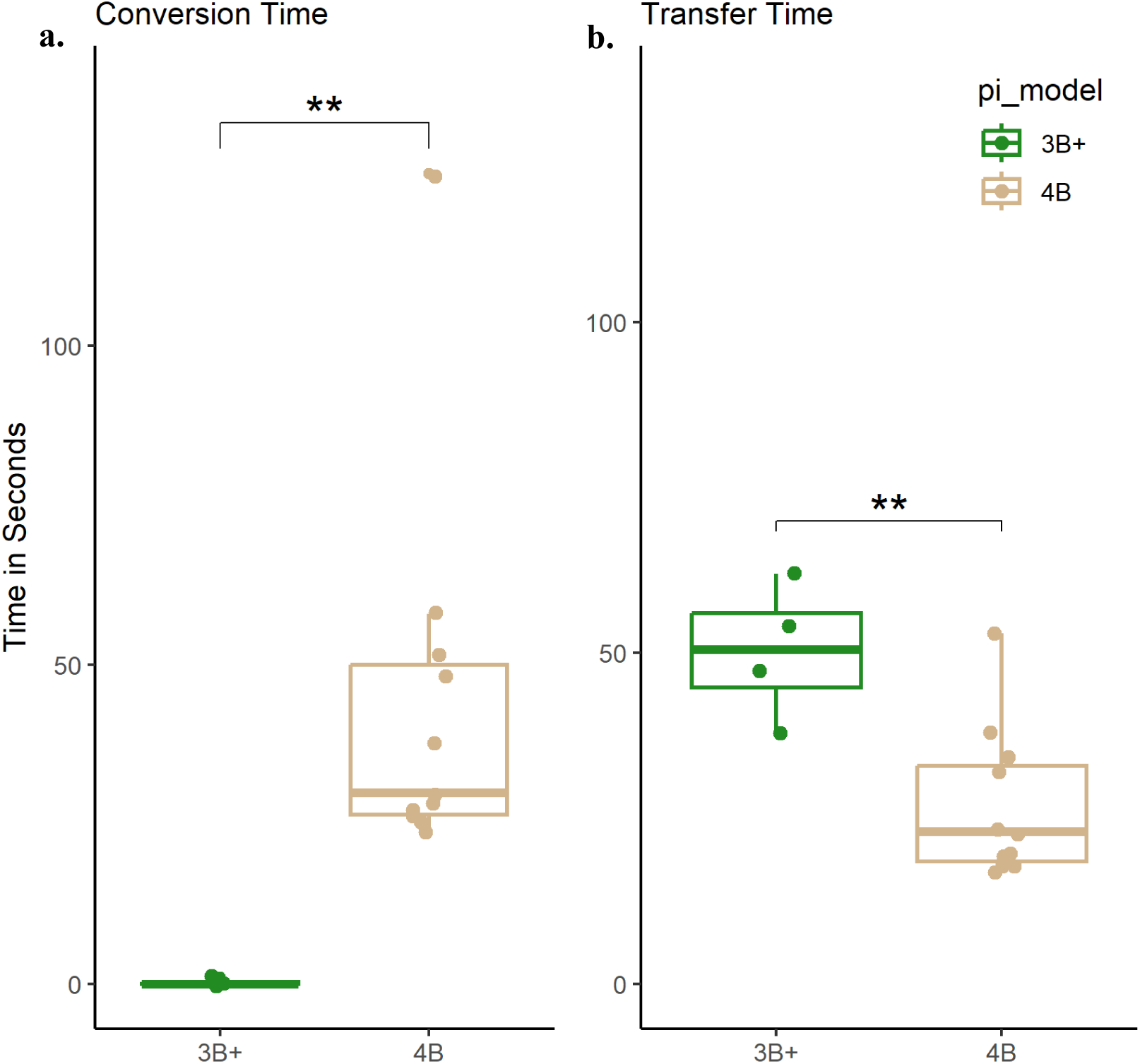
Boxplots of throughput comparisons between Raspberry Pi Models. Conversion Time (a.) and Transfer Time (b.).

##### 3.3.4.2. Fidelity

Analyses for the three full reference metric variables grouped by Pi Model revealed median scores of 0.99 ± 3e-4 (3B+) and 0.99 ± 4e-4 (4B) for SSIM, decibels of 44 ± 1dB (3B+) and 40 ± 0.3dB (4B) for PSNR, and 94 ± 0.3 (3B+) and 94 ± 0.2 (4B) for VMAF. Wilcoxon-Mann-Whitney tests revealed no significant differences in SSIM (*U*(4, 11) = 9, *p* = .104) or VMAF scores (*U*(4, 11) = 22, *p* = 1), but a significant difference was found in PSNR scores (*U*(4, 11) = 38, *p* = .04) with a large effect (*r* = 0.539) (Fig. 12.). Wilcoxon-Mann-Whitney tests found no significant differences between RPi models and IFIV (*U*(4, 11) = 27.5, *p* = .132) as only one recording (Cam3) from the 3B+ group experienced a single dropped frame.

**Fig. 12.**
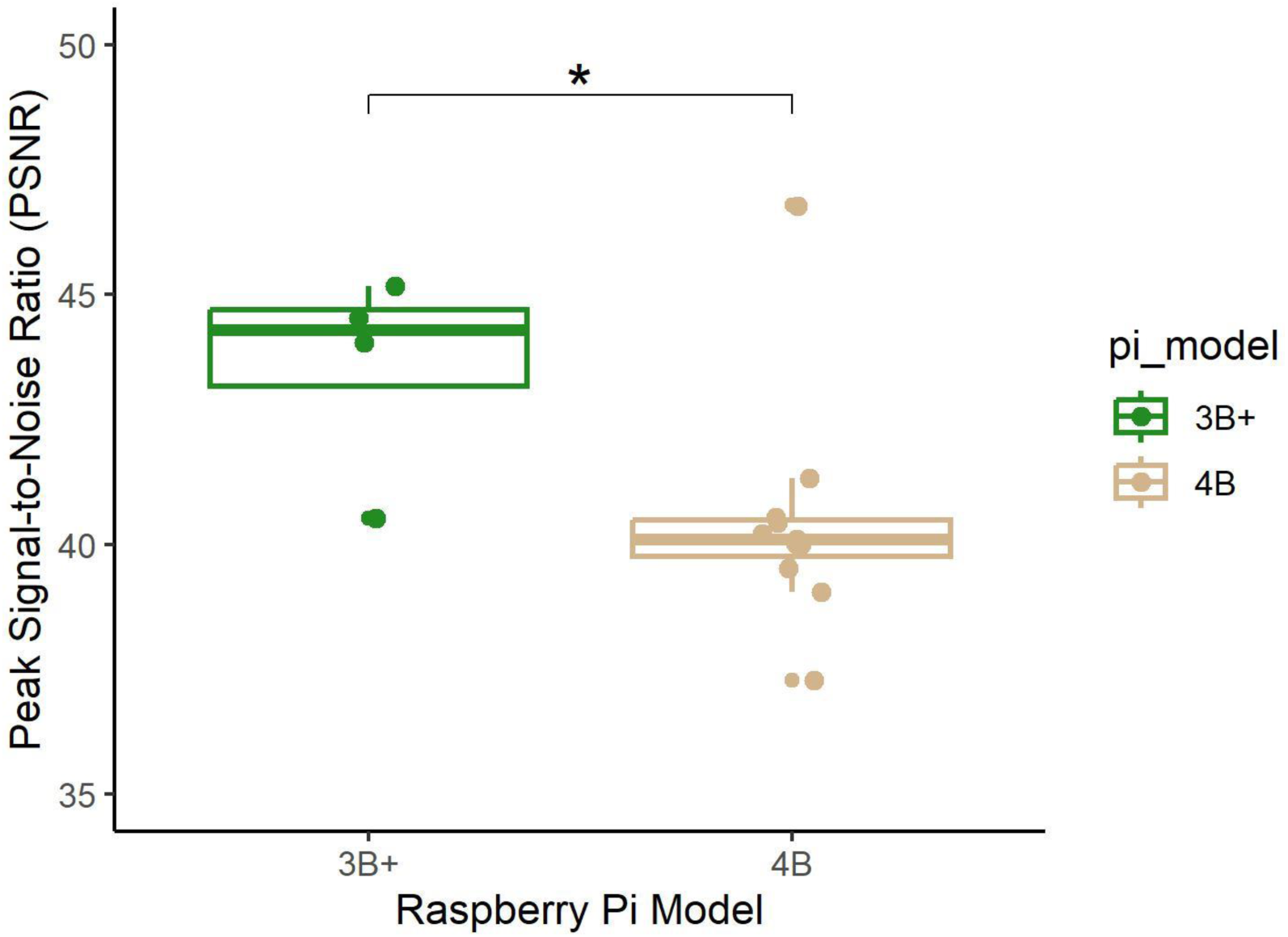
Peak Signal-to-Noise Ratio (PSNR). Boxplot visualizing significant differences in PSNR scores between Raspberry Pi models.

#### 3.3.5. Synchronicity

Fig. 13. below illustrates the statistical rationale to determine the light event frame and therefore time in seconds that the light event occurred as a measure of synchronicity across RPis in a given session. The full reference metric SSIM computes three separate metrics as a single score from -1 to 1, determining visual quality similarity between a reference image or video and a distorted (product) image or video, one of which is the luminance (Y) within an image.

**Fig. 13.**
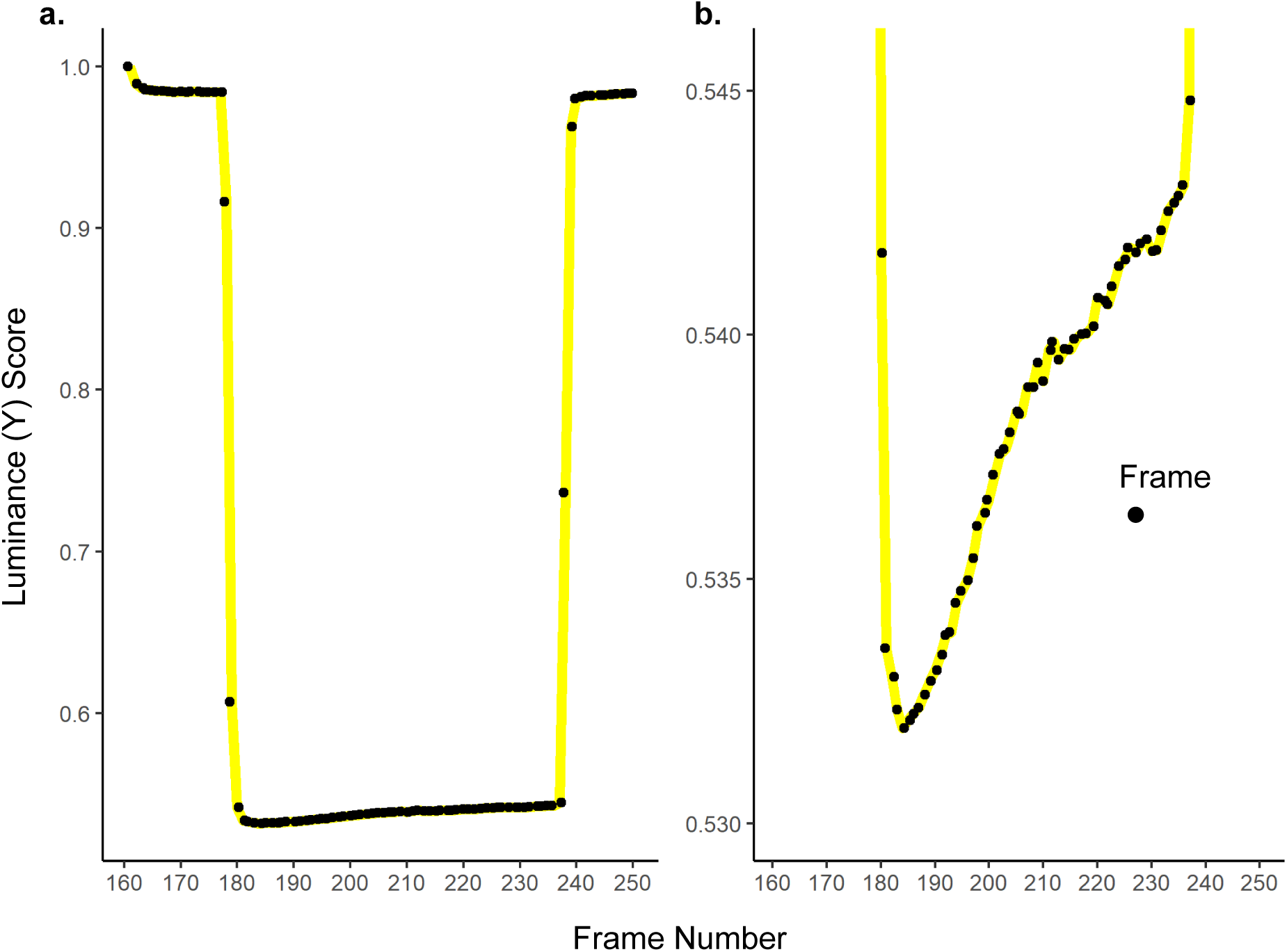
Light Event Identification. Line graphs showing Cam28 from session 2h-15 Boxes and each frame’s SSIM score to discover light event by inverse luminance (Y) score. **a.** The full set of individual frames were compared with reference frame 161, having a 1.0 after compared with its self, through to frame 250. **b.** Scaled to the frames in which the cue lights were on displaying the frame experiencing the brightest point (181) at the lowest Y value, and the increase in Y scores as the camera adjusts to the cue lights before frame 241 where cue lights turned off.

Therefore, if a dark image is used as the reference image of comparison, the frames, and thereby the particular frame in question, that the cue lights occur will have lower luminance scores given the dissimilarity in visual brightness. Figure 13. shows the results of one chamber’s SSIM scores across the full set of frames tested (13.a.) and a scaled view of the frames in which the cue lights were on (13.b.).

Once each frame was determined within a given recording, Levene’s tests of differences in variance by Time (*F*(1,38) = 0.03, *p* = .862) and by RPi models (*F*(1,13) = 1.36, *p* = .264), determined no significance differences in variances. For all recordings in the system (N=40), the mean light frame was 180 ± 0.7 or 6 ± 0.02 seconds.

### 3.4. Scalability

Commercial manufacturers such as Med Associates can only accommodate up to eight recording devices per computer. Many previous Pi-based studies have exhibited its scalability with 1-16 synchronized RPis in USB-type setups (Mathis et al., 2018; Nath et al., 2019; Hou and Glover, 2022), WiFi connections (Singh et al., 2019; Weber and Fisher, 2019, preprint, doi:10.1101/596106; Marcus et al., 2022), or through Ethernet (Saxena et al., 2018; Centanni and Smith, 2023). The rate at which one can acquire the necessary data or sample size for analyses is hindered by such low scales. Furthermore, many laboratories are housed in universities and technically secure locations, and a wirelessly connected RPi is a security risk to an institutional network. The use of RPis in conjunction with Ethernet switches allows for orders of magnitude more subjects per experiment. Additionally, the use of an ethernet switch removes the need to be directly connected to the institutional network, and subsequently results in the system being able to transmit significantly faster and be insensitive to the technical issues of the institutional network, meaning one can connect to the RPis despite internet failures.

### 3.5. Cost of Implementation

Commercial organizations, such as Med Associates, may provide state-of-the-art products, software, and functionality, but are expensive for an institution to implement. For instance, a single MedAssociates camera with a lens included costs ∼$779 (∼$46,740 for 60 cameras). Moreover, only eight cameras may be operated by a single computer, and only two computers can use a single software license, resulting in a cost of ∼$103,500 to record 60 chambers. Furthermore, the cameras utilized in Mathis et al. (2018) and Nath et al. (2019), the Point Grey made Firefly and Grasshopper3 4.1 MP Mono USB3 Vision (models: FMVU-03MTM-CS and CMOSIS CMV4000-3E12), Basler infrared-sensitive CMOS camera, Hero5 GoPro, and The Imaging Source made DFK-37BUX287 industrial camera, range from ∼$250 (Hero5 GoPro) to ∼$1,539 (Grasshopper3) for a single camera. Similarly, the Blackfly Mono S camera (model BFS-US-13Y3M-C) utilized in Pereira et al. (2022) are ∼$480 per camera as listed on Teledyne Flir.

The RPi is a frontrunner in addressing such issues of cost and use of space in a laboratory. For a 60-chamber system controlled by four computers, the cost of materials is approximately ∼$11,000 (Table. 1.), less than the camera products alone from Med Associates, and a fraction of the cost utilized in Mathis et al. (2018), Nath et al. (2019), and Pereira et al. (2022). Considering 23 RPis can be present on one Cisco Catalyst 3650 24-port switch, such an arrangement can be done across three computers at ∼$10,500, while a single fully stocked AWOW and Catalyst 24-port switch with 23 RPis equates to roughly ∼$4,000.

## 4. Discussion

This report demonstrates the fidelity, customizability, and highly scalable refinement of the affordable and high-throughput PiRATeMC (Pi-based Remote Acquisition Technology for Motion Capture) system with the synchronization of 60 Raspberry Pi video recordings for subsequent pose-estimation analysis. It was found that inclusion of the Arducam wide-angle fisheye lens provided better visibility of the full operant chamber and levers as opposed to the original camera lens. Development of the BoxTop device allows for precise installation, leading to greater stability and visibility across chambers and sessions than previous methods. Remote controllers and ethernet switchboards such as the AWOW mini-PC and Cisco Catalyst provide a localized network of instruments capable of initiating large-scale connections between devices.

The recording file *recordVideo.sh* is easily editable to fit the desired parameters of any laboratory, and capable of making multiple copies to fit specific experimental designs. Issues with embedded call variables were overcome by hard coding the password and video path.

Successful installation and execution resulted in recording 60 Raspberry Pis in parallel at considerably lower costs and video sizes, as well as higher flexibility than commercial alternatives with high video quality as shown in the results of SSIM, PSNR, VMAF, and IFIV. Analyses of throughput, fidelity, synchronicity, and comparisons of the most recent RPi models found that laboratories can virtually accomplish continuous data acquisition given the negligible processing time (*Mdn* = 116 ± 22 seconds) of recordings with quality assurance and synchronized within 0.02 seconds irrespective of RPi model.

The inclusion of the Arducam wide-angle fisheye lens provided better visibility of the full operant chamber and levers compared to the original camera lens. Experiments with large areas of operation, similar to that seen in Saxena et al. (2018), may benefit from wide-angle lenses by decreasing the number of cameras needed to capture the full experimental space given the Arducam is an 8MP (4K) device, consequently lowering costs without losing resolution.

Development of the BoxTop adapter allows for precision in installation, leading to greater stability and visibility across chambers than other methods in previous studies, particularly the use of hot glue in Singh et al. (2019), or Styrofoam holders outside the chambers in Weber and Fisher (2019, preprint, doi:10.1101/596106). Similarly, given Centanni and Smith (2023) report work with mice and therefore have smaller Med Associates provided chambers than the rat-sized chambers of this design, placement of the cameras 29cm from the grid floor results in full visibility of the chamber controls, but video is obscured by the top plane of glass as can be seen in the videos and images from their paper. This method would require regular maintenance to ensure clarity, while a lens such as the Arducam secured to the top pane of glass would capture the full area. Moreover, it is acknowledged that the 3D-created device by Hou and Glover (2022) is versatile and allows an extensive range of angle configurations and experimental designs. The use of 3D print was the original method of production for the BoxTop, but creation of a single device via 3D print was roughly 20 minutes, as opposed to no more than 1-hour to laser cut 150 devices from acrylic, allowing far greater numbers in a shorter time. A readily apparent limitation of the BoxTop is it only facilitates installation of cameras to the top of operant chambers and not for other forms of experiments such as open field or T-maze to name a few.

Remote controllers and ethernet switchboards such as the AWOW mini-PC and Cisco Catalyst featured in this system provide a headless network of instruments localized to a focal point capable of initiating a large-scale synchronicity between recording devices uncompromised by network issues. Mini-PCs are compact devices supplied in many hardware configurations to fit the needs and spatial requirements of any laboratory, while the Cisco Catalyst comes in varying numbers of ethernet ports with stackable capabilities leading to orders of magnitude more RPis to a single remote controller limited only by bandwidth.

The recording file *recordVideo.sh* is easily editable to fit the desired parameters of any laboratory, and capable of making multiple copies to fit specific experimental designs such as presented in this report. This customizability allows ease of execution and individuation of parameters, additional embedded codes or variables, video paths for storage, and inspiration for further coding development.

This system provides smaller video sizes at greater frame rates and comparative resolution than prior models that report such statistics. The video output results of Hou and Glover (2022) were 738.5MB per 29-minutes of video at 480p and 30FPS, Saxena (2019) resulted in 500MB (0.5GB) per hour of video at 640×480 resolution and 30FPS, while Weber and Fisher (2019, preprint, doi:10.1101/596106) experienced 440-500MB per hour during the light cycle and 1.5-2GB per hour in the dark cycle at 1280×768 15FPS. Comparatively, the present design video output results in only ∼350MB per hour at 800×600 resolution and 30FPS.

Descriptive statistics found that any given RPi (N=40) had a median conversion time of 55 ± 12.0 seconds and a median transfer time of 47.5 ± 10.6 seconds, for a total median processing time of 116 ± 22 seconds (∼2 minutes), displaying the capabilities of executing multiple sessions in a given day resulting in high video data output. If a lab were to run 1-hour sessions in succession and processing time invariably resulted in the high-end of the range (562 seconds), one could theoretically complete ∼20 hours of video data a day. Comparisons by Time revealed significant differences and large effects in conversion time (*U*(15, 25) = 36.5, *p* = < .001, *r* = 0.667), transfer time (*U*(15, 25) = 54.5, *p* = < .001, *r* = 0.588) and processing time (*U*(15, 25) = 38.5, *p* = < .001, *r* = 0.659), showing a scaling effect on length of session as processing time takes 2.7 times longer in a 2h session than a 1h session.

Tests of video fidelity confirmed the notion that the Raspberry Pi is not only a cost-efficient means of video data acquisition but one that does not sacrifice quality, as shown by the coefficient of variances of 0.19% (SSIM), 1.11% (VMAF), and 5.67% (PSNR), and only one RPi had a single dropped frame. Full reference metrics SSIM (0.99 ± 3e-4), VMAF (94 ± 0.2), and PSNR (40 ± 0.4), as well as IFIV (0.03 ± 3e-4) revealed high video quality of recordings, with each well within the range of desired scores. Furthermore, research shows SSIM and VMAF metrics are more representative to the human visual system (HVS) and often outperforms PSNR which is also vulnerable to manipulations (Vranješ et al., 2013; Li, et al., 2016; Deshpande, Ragha, & Sharma, 2018; Li et al., 2018; Sara, Ackter, & Uddin, 2019).

Variance analyses confirmed synchronicity as the system obtained a light event frame standard deviation of 4.32 frames (*M* = 180 ± 1) or 144 milliseconds (*M* = 6 ± 0.02 seconds). This displays accuracy of the RPi and its potential use in timed tasks and systems requiring temporal precision.

Comparisons of the latest Raspberry Pi models 3B+ and 4B found significant differences in conversion time and transfer time, but not in processing time, as well as significant differences in PSNR scores. The differences in conversion time and transfer time negate one another in light of the full processing time. Although PSNR scores differ by as much as ∼7dB, SSIM and VMAF scores are virtually identical between groups. These differences, although significant, have negligible effects on a laboratory’s pursuits and regress to personal preference or market availability of the 3B+ or 4B models.

Lastly, a major advantage of the RPi is its affordability and versatility to suit the needs of any laboratory as a number of configurations can be made with the removal or addition of other items. To fully place the RPi in perspective within the discussion of recording device options, a single fully stocked Raspberry Pi, complete with RPi, camera, fish-eye lens, 18” cable, PoE+ HAT, infrared light and adapter, micro-SD card, and Ethernet cable, cost around ∼$150, and with computer and switchboard included is ∼$670, considerably less than commercial alternatives.

Therefore, the cost of implementation ranges from $670 (1 RPi) to ∼$11,000 (60 RPis) as outlined in this system. Furthermore, the Raspberry Pi utilizes the elements present on the Linux/Ubuntu OS to conduct its functions, eliminating the need for new software purchases or installations, and capable with a few lines of code in the terminal to establish an interface between the remote and switch.

## 5. Conclusion

This report demonstrates an affordable, customizable, and highly scalable implementation and expansion of the PiRATeMC (Pi-based Remote Acquisition Technology for Motion Capture) system with the synchronization of 60 Raspberry Pi video recordings for subsequent pose-estimation analysis. A laboratory can be fully installed with minimal coding experience in a month’s time by a single technician, providing the ability to record a large number of subjects in a variety of species and settings, resulting in a high throughput and fidelity of video data with low storage needs and financial burdens. The Raspberry Pi’s versatility and affordability makes it an attractive option for various laboratory needs and designs at a competitive price compared to commercial products with no tradeoff of video quality or temporal precision.

## 6. CRediT authorship contribution statement

**Jarryd Ramborger:** Conceptualization, Data curation, Formal analysis, Investigation, Methodology, Project administration, Software, Visualization, Writing – original draft. **Alexander C.W. Smith, PhD:** Conceptualization, Methodology, Software, Validation, Writing – review & editing. **Olivier George, PhD:** Conceptualization, Data curation, Funding acquisition, Investigation, Methodology, Project administration, Resources, Supervision, Validation, Writing – review & editing. **Sumay Kalra:** Data curation, Investigation, Project administration, Software.

## 7. Code Availability

Coding and tutorial to replicate this system found on GitHub at: https://github.com/George-LabX/raspicluster/blob/main/README.md. Coding and tutorial for the PiRATeMC system found on GitHub at: https://github.com/alexcwsmith/PiRATeMC/blob/master/README.md. This project makes use of the FFmpeg library, a leading multimedia framework that provides a comprehensive suite of tools for processing audio, video, and other multimedia content, version 2023-08-17-git-9ae4863cc5-full_build-www.gyan.dev, developed and copyrighted by the FFmpeg developers between 2000 and 2023, and compiled with gcc 12.2.0 (Rev10) by the MSYS2 project. FFmpeg source code can be accessed at https://ffmpeg.org/download.html. FFmpeg and Windows PowerShell codes for analyses, as well as R Studio R Markdown can be accessed at https://github.com/George-LabX/raspicluster/tree/main/RPi_Codes/RPi_paper_analyses

## Acknowledgements

Authors would like to thank and acknowledge Dr. Centanni and Dr. Smith for allowing their Raspberry Pi design to be open source on GitHub for laboratories to utilize and customize for the advancement of research per the BSD 3-Clause License.

Copyright (c) 2023, Alexander CW Smith

Redistribution and use in source and binary forms, with or without modification, are permitted provided that the following conditions are met:

1. Redistributions of source code must retain the above copyright notice, this list of conditions and the following disclaimer.
2. Redistributions in binary form must reproduce the above copyright notice, this list of conditions and the following disclaimer in the documentation and/or other materials provided with the distribution.
3. Neither the name of the copyright holder nor the names of its contributors may be used to endorse or promote products derived from this software without specific prior written permission.

THIS SOFTWARE IS PROVIDED BY THE COPYRIGHT HOLDERS AND CONTRIBUTORS “AS IS” AND ANY EXPRESS OR IMPLIED WARRANTIES, INCLUDING, BUT NOT LIMITED TO, THE IMPLIED WARRANTIES OF MERCHANTABILITY AND FITNESS FOR A PARTICULAR PURPOSE ARE DISCLAIMED. IN NO EVENT SHALL THE COPYRIGHT HOLDER OR CONTRIBUTORS BE LIABLE FOR ANY DIRECT, INDIRECT, INCIDENTAL, SPECIAL, EXEMPLARY, OR CONSEQUENTIAL DAMAGES (INCLUDING, BUT NOT LIMITED TO, PROCUREMENT OF SUBSTITUTE GOODS OR SERVICES; LOSS OF USE, DATA, OR PROFITS; OR BUSINESS INTERRUPTION) HOWEVER CAUSED AND ON ANY THEORY OF LIABILITY, WHETHER IN CONTRACT, STRICT LIABILITY, OR TORT (INCLUDING NEGLIGENCE OR OTHERWISE) ARISING IN ANY WAY OUT OF THE USE OF THIS SOFTWARE, EVEN IF ADVISED OF THE POSSIBILITY OF SUCH DAMAGE.

The licensing terms for FFmpeg are governed by the GNU Lesser General Public License version 2.1 (LGPLv2.1)(http://www.gnu.org/licenses/old-licenses/lgpl-2.1.html). For further information and access to FFmpeg’s features and documentation, please visit the official https://ffmpeg.org website. Adhering to the LGPLv2.1 license conditions is imperative when utilizing or disseminating the combined work.

## 10. Declaration of Generative AI and AI-assisted technologies in the writing process

*Statement: During the preparation of this work the author(s) used ChatGPT-4 in order to accelerate coding for FFmpeg analyses. After using this tool/service, the author(s) reviewed and edited the content as needed and take(s) full responsibility for the content of the publication*.

## References

1. A. K. Venkataramanan, C. Wu, A. C. Bovik, I. Katsavounidis and Z. Shahid, “A Hitchhiker’s Guide to Structural Similarity,” in IEEE Access, vol. 9, pp. 28872–28896, 2021, doi: 10.1109/ACCESS.2021.3056504.

(2) Basler infrared-sensitive CMOS camera for purchase on Direct Industry. https://www.directindustry.com/product-manufacturer/basler-infrared-camera-73401-3007.html (last accessed 15 July 2023).

(3) Blackfly S 13YM3-M USB3 camera for purchase on Teledyne Flir. https://www.flir.com/products/blackfly-s-usb3/?model=BFS-U3-13Y3M-C&vertical=machine+vision&segment=iis (last accessed 19 July 2023).

4. Belin, D., Balado, E., Piazza, P. V., & Deroche-Gamonet, V. (2009). Pattern of intake and drug craving predict the development of cocaine addiction-like behavior in rats. Biological Psychiatry, 65(10), 863–868.

5. Centanni, S.W., Smith, A.C.W. (2023). PiRATeMC: A highly flexible, scalable, and low-cost system for obtaining high quality video recordings for behavioral neuroscience. Addiction Neuroscience, 8, 100108. ISSN 2772-3925. 10.1016/j.addicn.2023.100108.

6. Deshpande, R. G., Ragha, L. L., & Sharma, S. (2018). Video Quality Assessment through PSNR Estimation for Different Compression Standards. Indonesian Journal of Electrical Engineering and Computer Science, 11(3), 918–924. DOI:10.11591/ijeecs.v11.i3.pp918-924

(7) FFmpeg website https://ffmpeg.org/ffmpeg.html (last accessed 04 September 2023).

(8) George Lab GitHub instructions and tutorial. https://github.com/George-LabX/raspicluster/blob/main/README.md (last accessed 27 May 2023).

9. Guglielmo, D., et al. (2023). Characterization of cocaine addiction-like behavior in heterogeneous stock rats. bioRxiv. 10.1101/2021.07.22.453410.

(10) Hero5 GoPro for purchase on Amazon. https://www.amazon.com/GoPro-HERO5-Black-Waterproof-Digital/dp/B01M14ATO0 (last accessed 16 July 2023).

11. Hou, S., Glover, E.J. (2022). Pi USB Cam: A Simple and Affordable DIY Solution That Enables High-Quality, High-Throughput Video Capture for Behavioral Neuroscience Research. eNeuro 7 September 2022, 9 (5) ENEURO.0224-22.2022; DOI: 10.1523/ENEURO.0224-22.2022.

(12) The Imaging Source made DFK-37BUX287 industrial camera for purchase on Wilco Imaging. https://wilcoimaging.com/products/dfk-37bux287 (last accessed 16 July 2023).

13. Kallupi, M., de Guglielmo, G., Carrette, L. L. G., Simpson, S., Kononoff, J., Kimbrough, A., Smith, L. C., Shankar, K., Avelar, A., Conlisk, D., Brennan, M., Tieu, L., Sedighim, S., Boomhower, B., Maturin, L., Fannon, M. J., Martinez, A., Crook, C., Dirik, S., Velarde, N., Schweitzer, P., Bonnet-Zahedi, S., Othman, D. N., Sichel, B., Guess, K., Peng, B., Hu, A. S., Chun, L. E., Milan, K., Lau, J., Zheng, Y., Vang, A., Solberg Woods, L. C., Palmer, A. A., & George, O. (2022). Individual differences in oxycodone addiction-like behaviors in a large cohort of heterogeneous stock (HS) rats. bioRxiv, 2022.07.26.501618. 10.1101/2022.07.26.501618.

(14) Li, Z., Aaron, A., Katsavounidis, I., Moorthy, A., & Manohara, M. (2016, June 6). Toward A Practical Perceptual Video Quality Metric. Netflix TechBlog. https://netflixtechblog.com/toward-a-practical-perceptual-video-quality-metric-653f208b9652.

(15) Li, Z., Bampis, C., Novak, J., Aaron, A., Swanson, K., Moorthy, A., & De Cock, J. (2018, October 25). VMAF: The Journey Continues. Netflix TechBlog. https://netflixtechblog.com/vmaf-the-journey-continues-44b51ee9ed12

16. Luxem, K., Mocellin, P., Fuhrmann, F. et al. Identifying behavioral structure from deep variational embeddings of animal motion. Commun Biol 5, 1267 (2022). 10.1038/s42003-022-04080-7.

17. Marcus, A. D., Achanta, S., & Jordt, S. E. (2022). Protocol for non-invasive assessment of spontaneous movements of group-housed animals using remote video monitoring. STAR Protocols, 3(2), 101326. 10.1016/j.xpro.2022.101326.

18. Mathis, A., Mamidanna, P., Cury, K.M. et al. DeepLabCut: markerless pose estimation of user-defined body parts with deep learning. Nat Neurosci 21, 1281–1289 (2018). 10.1038/s41593-018-0209-y.

19. Nath, T., Mathis, A., Chen, A.C. et al. Using DeepLabCut for 3D markerless pose estimation across species and behaviors. Nat Protoc 14, 2152–2176 (2019). 10.1038/s41596-019-0176-0.

(20) PiRATeMC (Pi-based Remote Acquisition Technology for Motion Capture) GitHub tutorial and instructions. https://github.com/alexcwsmith/PiRATeMC/blob/master/README.md last (accessed 14 July 2023).

(21) Point Grey Firefly (model FMVU-03MTM-CS) for purchase on Radwell. https://www.radwell.com/en-US/Buy/POINT%20GREY/POINT%20GREY/FMVU-03MTM-CS (last accessed 15 July 2023).

(22) Point Grey Grasshopper3 4.1 MP Mono USB3 Vision (model CMOSIS CMV4000-3E12) for purchase on Teledyne Flir. https://www.flir.com/products/grasshopper3-usb3/?model=GS3-U3-41C6NIR-C&vertical=machine+vision&segment=iis (last accessed 15 July 2023).

(23) PowerShell coding and syntax assistance: https://learn.microsoft.com/en-us/powershell/module/microsoft.powershell.management/?view=powershell-7.3 (last accessed 04 September 2023).

24. Sara, U., Akter, M. and Uddin, M. (2019) Image Quality Assessment through FSIM, SSIM, MSE and PSNR—A Comparative Study. Journal of Computer and Communications, 7, 8–18. doi: 10.4236/jcc.2019.73002.

25. Saxena R, Barde W, Deshmukh SS (2018) Inexpensive, scalable camera system for tracking rats in large spaces. Journal of Neurophysiology 120:2383–2395.

(26) Singh S, Bermudez-Contreras E, Nazari M, Sutherland RJ, Mohajerani MH (2019) Low-cost solution for rodent home-cage behaviour monitoring Silvani A, ed. PLoS ONE 14:e0220751.

27. Spanagel, R. (2017). Animal models of addiction. Dialogues in Clinical Neuroscience, 19(3), 247–258. doi: 10.31887/DCNS.2017.19.3/rspanagel PMID: 29302222; PMCID: PMC5741108.

28. Setiadi, D.I.M (2021). PSNR vs SSIM: imperceptibility quality assessment for image steganography. Multimed Tools Appl 80, 8423–8444. 10.1007/s11042-020-10035-z

29. T.D. Pereira, N. Tabris, A. Matsliah, D. M. Turner, J. Li, S. Ravindranath, E. S. Papadoyannis, E. Normand, D. S. Deutsch, Z. Y. Wang, G. C. McKenzie-Smith, C. C. Mitelut, M. D. Castro, J. D’Uva, M. Kislin, D. H. Sanes, S. D. Kocher, S. S-H, A. L. Falkner, J. W. Shaevitz, and M. Murthy. Sleap: A deep learning system for multi-animal pose tracking. Nature Methods, 19(4), 2022.

30. Vranješ, M., Rimac-Drlje, S., & Grgić, K. (2013). Review of objective video quality metrics and performance comparison using different databases. Signal Processing: Image Communication, 28(1), 1–19. ISSN 0923-5965. 10.1016/j.image.2012.10.003.

31. Wang Z, Bovik AC, Sheikh HR, Simoncelli EP. Image quality assessment: from error visibility to structural similarity. IEEE Trans Image Process. 2004 Apr;13(4):600–12. doi: 10.1109/tip.2003.819861. PMID: 15376593.

(32) Weber WD, Fisher HS (2019) An inexpensive remotely operated video recording system for continuous behavioral observations. Animal Behavior and Cognition. http://biorxiv.org/lookup/doi/10.1101/596106.

